# Deep Learning-Based Spatial Immunoprofiling of Multiplex Immunofluorescence Images Distinguishes Tuberculosis Disease States in Diversity Outbred Mice

**DOI:** 10.64898/2026.01.26.701667

**Authors:** Usama Sajjad, Nicholas A. Crossland, Metin N. Gurcan, Gillian Beamer, Muhammad Khalid Khan Niazi

**Affiliations:** Department of Pathology, The Ohio State University Wexner Medical Center, Columbus, OH, USA; National Emerging Infectious Diseases Laboratories (NEIDL), Boston University, Boston, MA, USA; Department of Pathology and Laboratory Medicine, Boston University Chobanian & Avedisian School of Medicine, Boston, MA, USA; Department of Virology, Immunology, & Microbiology, Boston University Chobanian & Avedisian School of Medicine, Boston, MA, USA; Center for Artificial Intelligence Research, Wake Forest University School of Medicine, Winston-Salem, NC, USA; Texas Biomedical Research Institute, San Antonio, Texas, USA

## Abstract

Tuberculosis (TB), caused by *Mycobacterium tuberculosis* (*M*.*tb*), remains a major global health challenge, with approximately 10.8 million new cases and 1.25 million deaths reported in 2023. Human responses to *M*.*tb* are heterogeneous with clinical outcomes including bacterial clearance, asymptomatic latent *M*.*tb* infection, and severe fatal pulmonary TB. Here, we aim to address knowledge gaps in the organization of *M*.*tb* granulomas by identifying cell-based spatial features indicating asymptomatic lung infection. To address this gap, which cannot be directly studied in humans, we used lung sections from *M*.*tb* infected Diversity Outbred mice with acute pulmonary TB, asymptomatic *M*.*tb* infection, or chronic pulmonary TB that were stained for T cells, B cells, macrophages, and bronchiolar epithelial cells by multiplexed immunofluorescence. We first developed a new, accurate model to automatically segment lung granulomas, detect/quantify the cell types within granulomas, and extract the location of immune cells in granulomas for each disease state. Analysis of model-derived results show that lung granulomas from asymptomatic mice have a characteristic spatial profile consisting of higher CD4+ and CD8a+ T cell densities, closer B cell proximity to bronchiolar epithelium, and increased T cell-macrophage proximity in asymptomatic *M*.*tb* infection. Next, we propose a second novel approach to utilize a large language model (LLM) to independently decode complex cellular patterns within granulomas, and distinguished key immunological signatures: balanced immune expression in asymptomatic *M*.*tb* mice, dysfunctional responses with low cellularity in acute TB, and highest immune cells in chronic TB. Overall, the results from this study show that lung granulomas in asymptomatic infection are characterized by increased T cell density, increased numbers of peribronchiolar B cells, and T cells closer to macrophages as compared to acute and chronic TB. These methods and results help establish an automated pipeline to extract and analyze data from multiplexed fluorescence images and provide a foundation to better understand how granuloma architecture varies by disease state.

## Introduction

Tuberculosis (TB), an infectious disease caused by the bacterial pathogen *Mycobacterium tuberculosis* (*M*.*tb*), remains a substantial global health threat. Epidemiological studies estimate that about one-quarter of the global population has been infected with *M*.*tb* (1). In 2023, the global burden of TB reached 10.8 million new cases, increased from 10.1 million cases in 2020 (1). In adult humans, infection with *M*.*tb* results a clinical spectrum including acute fulminant pulmonary TB with rapid progression to death within weeks; chronic pulmonary TB with slower lung-damaging progression often lasting years; and asymptomatic infection (e.g., incipient TB, latent *M*.*tb* infection (LTBI)); and clearance (2, 3). Symptoms of pulmonary TB are cough, fever, weight loss, and cachexia, due to tissue-destructive inflammation (4). Notably, most humans with asymptomatic LTBI who progress to active pulmonary TB do so in the absence of identifiable risk factors (5), underscoring the multifactorial determinants of pulmonary TB progression, unpredictable nature of reactivation, and the need to improve our understanding of granulomas in different disease states, including asymptomatic LTBI.

*M*.*tb* infection initiates granuloma formation (6), which is the hallmark of infection that develops in response to molecular and cellular interactions between immune cells and bacterial components (7). Granuloma formation is believed to begin with activation, recruitment and aggregation of leukocytes surrounding *M*.*tb*-infected macrophages, creating a protective barrier that limits bacterial dissemination (6), restricts *M*.*tb* replication and maintains LTBI. The spatial organization of immune cells within granulomas across different disease states, particularly the proximity of immune cells, macrophages, T cells, B cells, and bronchiolar epithelial cells and how these spatial relationships differ between asymptomatic LTBI, acute pulmonary TB, and chronic active TB are not fully known. Since controlled *M*.*tb* lung infection experiments cannot be directly performed in humans, we used lung sections from *M*.*tb-*infected DO mice. Each DO mouse represents a heterozygous mosaic of DNA inherited from eight founder strains (8, 9), and when infected with *M*.*tb*, DO mice recapitulate human TB pathology more accurately than inbred strains of mice (10, 11). Here, we used multiplexed immunofluorescence imaging (mIF) of lung tissue sections from DO mice, to study and visualize the spatial location of CD19+ B cells, CD4+ helper T cells, CD8a+ cytotoxic T cells, EpCAM+ epithelial cells, and IBA1+ macrophages.

Given the structural complexity of granulomas, manual analysis cannot capture the spatially relevant relationships amongst immune cells and epithelial cells. The transition from traditional histopathology examination of glass slides to whole-slide imaging (WSI), combined with advances in high-speed data transfer and cost-effective storage, has enabled deep learning-based identification of complex histopathological patterns and feature quantification that is not possible by visual examination (12-17). This advancement supported several new discoveries, including our prior work on imaging biomarkers in lung granulomas of *M*.*tb*-infected mice with and without BCG vaccination, and accurate prediction of gene expression based on granulomas\s (2, 14, 16, 18).

Despite recent progress in applying deep learning to hematoxylin and eosin (H&E)–stained whole-slide images for tuberculosis phenotyping, these approaches remain biologically limited. H&E-based models primarily rely on global morphological pattern recognition, thereby obscuring cellular specificity and spatial immune interactions that are central to tuberculosis pathogenesis. Critical biological processes, including immune cell recruitment, compartmentalization, and cell–cell proximity within granulomas, cannot be resolved from H&E staining alone, as immune cell subtypes and functional states are visually indistinguishable. As a result, while H&E-based deep learning models can achieve disease-state classification, they provide less insight into the immune cell types that contribute to acute pulmonary TB, asymptomatic infection, and chronic pulmonary TB in DO mice. In contrast, mIF imaging enables simultaneous, cell-type–specific visualization of immune and epithelial populations and allows explicit quantification of spatial relationships within granulomas.

Previous studies in humans with pulmonary TB have shown that granuloma necrosis is at the center of granulomas, and is a distinguishing feature DO mice with acute pulmonary TB (19-22), while an abundance of B-cells in perivascular and peribronchiolar cuffs identify asymptomatic infection in DO mice (3). The spatial relationship between immune cells and bronchiolar epithelial cells was a key motivation for using mIF imaging, yet manual quantification of these spatial patterns across entire lung sections was intractable. To address this limitation, we present an end-to-end framework that automates granuloma segmentation, classifies TB states more accurately than previous H&E-based methods, and quantifies spatial relationships between immune cell populations in granulomas. We replicate and validate the peribronchiolar B-cell signature of asymptomatic infection (3) using an independent method. By measuring the distance between CD19+ B-cells and bronchiolar epithelial cells in mIF images, we show that asymptomatic mice exhibit significantly closer B-cell-epithelial cell proximity than DO mice with acute pulmonary TB (“progressors” that survive less than 60 days) or chronic pulmonary TB (“controllers” that survive greater than 60 days), independently confirming previous findings (3) using automated spatial analysis. Finally, we integrate large language models to interpret complex cellular patterns by synthesizing cell identities, spatial coordinates, expression intensities, and attention-derived importance scores, extracting biologically interpretable insights that distinguish TB disease states. This approach also addresses the limitations of traditional hypothesis-driven methods, which, while valuable, can be constrained by fixed assumptions and human bias, potentially overlooking the complex, multi-dimensional biological patterns that emerge from integrating multi-modal data. By identifying relationships across these diverse data modalities without predetermined hypotheses, LLMs enable discovery of novel biological insights that may be difficult to capture through conventional statistical methods alone. Overall, this is the first study to combine mIF, attention-based MIL, and LLM-driven interpretation to distinguish TB disease states at the level of the granuloma in a genetically diverse animal model.

## Data

### 2.1. Ethics Statement

All lung tissue sections used in this study were generated from experiments conducted under protocols approved by the Institutional Animal Care and Use Committee (IACUC) at Tufts University (protocol numbers: G2012-53, G2015-33, G2018-33, and G2020-121). Additionally, the Institutional Biosafety Committee (IBC) at Tufts University approved all experimental procedures involving *M*.*tb* under registration numbers GRIA04, GRIA10, GRIA17, and 2020-G61. All animal handling and experimental procedures adhered to institutional guidelines for the ethical treatment of laboratory animals.

### 2.2. Experimental Animals and Tissue Samples

DO mice (J:DO strain 009376, RRID:IMSR_JAX:009376) from generations 20 to 32 were obtained from The Jackson Laboratory (Bar Harbor, ME) at 4 weeks of age. At 8 to 10 weeks of age, all mice were exposed to a low-dose aerosol infection of *M*.*tb* as previously described (3, 23). This infection method ensures a physiologically relevant infectious dose within the predicted infectious dose for humans (24), and DO mice develop a spectrum of pulmonary TB that recapitulates some of the disease states observed in humans (3).

Our dataset comprises 48 mIF lung tissue sections from experimentally infected DO mice representing three distinct disease states: 15 mice with Progressors, 17 mice classified as Asymptomatic, and 16 mice as Controllers. These disease classifications were determined based on established clinical and histopathological criteria as described in previous studies (3, 25)

## Methods

Previous studies have established that granuloma necrosis is at the center of granulomas, and a distinguishing feature of super susceptible progressor DO mice (19-22), while B-cells in perivascular and peribronchiolar cuffs identify asymptomatic latent infection (3). The spatial relationship between B-cells and bronchiolar epithelial cells was a key motivation for developing mIF imaging, yet manual quantification of these spatial patterns across entire lung sections remains intractable. To address this limitation, we present an end-to-end framework that automates granuloma segmentation, classifies disease states with substantial improvements over H&E-based methods, and quantifies spatial relationships between immune cell populations at the granuloma level. In this section, we discuss the methodology of our framework in detail. Our dataset consists of entire lung sections, requiring an initial step to segment the TB granulomas. From the segmented images, we then train a deep learning model to predict disease state. Finally, we use an LLM to interpret the spatial patterns of immune cells within high-attention regions identified by our classification model, extracting biologically meaningful insights that distinguish the three disease states. The entire workflow of our framework is presented in Figure 1.

**Figure 1.**
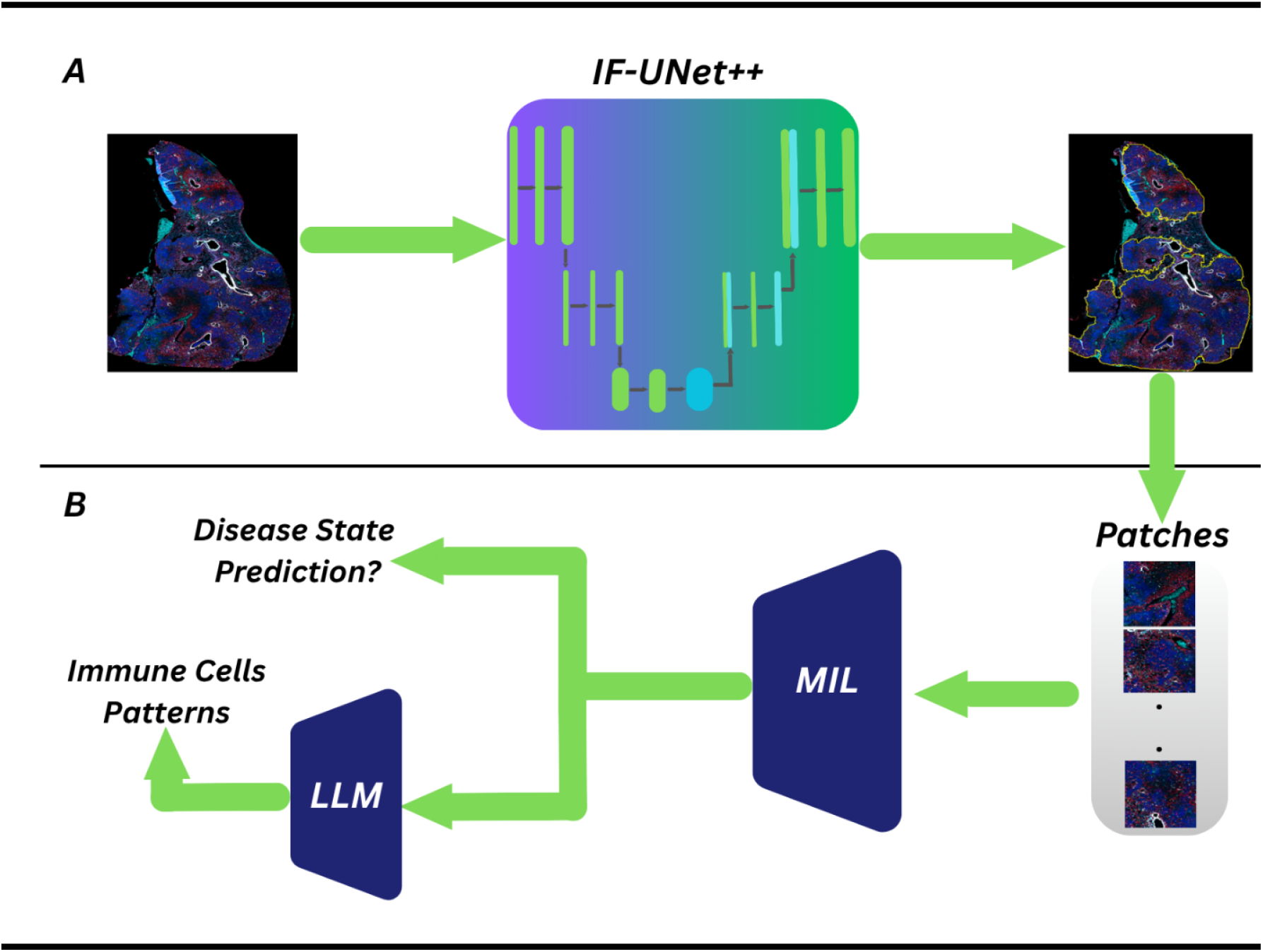
An overview of our proposed framework. **A**. mIF images are processed using the IF-UNet++ model to segment granulomas. The resulting segmentation outputs are divided into patches for further analysis. **B**. These patches are fed into a MIL model to predict disease-state classification. A trained MIL model, along with an LLM, integrates MIL-derived features and immune cell spatial patterns to infer immune cell patterns.

### 3.1. Segmentation of Granulomas from Lung Tissue Sections

The objective of the segmentation component is to accurately delineate tuberculosis granulomas from multiplex immunofluorescence whole-slide images, thereby defining biologically meaningful regions for downstream spatial and disease-state analysis. Each lung section contains complex and heterogeneous tissue architecture, necessitating an automated approach capable of capturing granuloma boundaries across diverse immune phenotypes. To this end, we employ an immunofluorescence-adapted UNet++ architecture (IF-UNet++), which leverages multiscale feature aggregation and dense skip connections to preserve fine-grained cellular and structural information across multiple immunofluorescence channels.

To study the granulomas, we first identify granulomas from mouse lung WSIs. Given *N* WSIs, we extract *M* non-overlapping patches, where *M* may vary per WSI depending on its size. Each input patch *x*_*i*_ ∈ ℝ ^*H* × *W* × *D*^ comprises *H* × *W* pixels across *D* immunofluorescence channels, with each channel corresponding to a distinct cellular marker. The corresponding ground truth segmentation mask *y*_*i*_ ∈ {0,1}^*H* × *W*^ represents a binary mask delineating the granuloma boundary. This yields a dataset 𝒟 comprising of paired patches (*x*_*i*_) and its ground truth (*y*_*i*_). Our objective is to learn a mapping function *f*: *x* → *y* that accurately predicts the segmentation mask *y*_*i*_ for each input patch *x*_*i*_.

We propose an IF-UNet++ architecture (26) optimized for immunofluorescence image segmentation to learn this mapping function *f*. Our architecture employs a nested encoder-decoder structure interconnected through dense skip pathways (26). The encoder performs progressive downsampling while computing hierarchical feature representations at multiple spatial resolutions. Unlike the standard U-Net, which directly concatenates encoder features with decoder features at matching resolutions, IF-UNet++ introduces intermediate convolution blocks along each skip pathway. These blocks iteratively refine encoder representations before fusion with decoder outputs. At each computational node, we aggregate features from all preceding nodes along the pathway and upsample features from deeper layers through a series of convolution operations, enabling dense information flow across network hierarchies.

We train our network with deep supervision, attaching segmentation heads at multiple semantic levels. The loss function combines binary cross-entropy and Dice coefficient, enabling the model to learn representations at different scales simultaneously. During inference, the final segmentation mask is generated by averaging outputs from all segmentation branches, producing pixel-wise predictions that identify granuloma regions within each patch. The overall loss function for the segmentation model is defined as (Eq. 1):

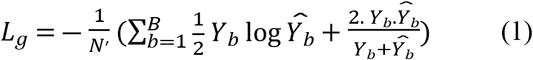

where *B* represents the batch size, *Y*_*b*_ denotes the flattened ground truth segmentation mask for the *b*^*th*^ image, and 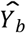 represents the flattened predicted probability map. This formulation combines two complementary objectives to optimize segmentation performance. The first term represents binary cross-entropy, which measures pixel-wise classification error and encourages accurate probability estimates at each pixel location. The second term corresponds to the Dice coefficient, which quantifies overlap between predicted and ground truth masks. The negative sign converts this into a minimization problem. By minimizing *L*_*g*_ during training, the network learns to generate accurate pixel-wise predictions that effectively segment granulomas from lung tissue sections.

During WSI-level inference, we partition incoming WSI into non-overlapping patches, process each patch independently using *f*, and reconstruct the full segmentation map by spatially stitching the predicted outputs to generate a complete WSI-level segmentation.

### 3.2. Multiple Instance Learning for Disease State Classification

Whole-slide lung sections from M. tuberculosis–infected mice exhibit substantial intra-slide heterogeneity, with multiple granulomas and surrounding tissue regions contributing differently to disease phenotype. Assigning patch-level disease labels is therefore biologically ill-defined, as disease state emerges from the collective behavior of multiple granulomas rather than from any single localized region. Multiple instance learning (MIL) provides a principled framework for this setting by enabling slide-level classification while allowing the model to learn which granuloma regions are most informative for disease discrimination (27-29). Importantly, attention-based MIL further permits identification of high-impact regions, offering a direct link between model predictions and biologically meaningful tissue compartments.

In this section, we describe the Multiple Instance Learning (MIL) model (30) used to classify lung tissue sections from *M*.*tb-*infected DO mice into three distinct disease states: asymptomatic infection, acute pulmonary tuberculosis in progressors, and chronic pulmonary tuberculosis in controllers.

Our training set consists of *N* gigapixel-scale WSIs, where each WSI (*X*_*i*_) represents a DO mouse lung tissue section containing one or more granulomas. These WSIs are denoted as bags:

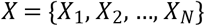

with corresponding disease-state labels

*Y* = {*Y*_1_, *Y*_2_, …, *Y*_*N*_}where *Y*_*l*_ ∈ {0, 1, 2} for *l* ∈ {1,…, *N*}, corresponds respectively to asymptomatic, acute, and chronic TB states.

Under the MIL formulation, each WSI is decomposed into a set of non-overlapping image patches (instances),

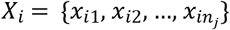

where *n*_*j*_ denotes the total number of patches extracted from the *i*^*th*^ WSI, and *j* = 1,…,*n*_*j*_ indexes the patches within the *i*^*th*^ WSI. Each patch *x*_*ij*_ ∈ ℝ ^*H* × *W* × *C*^ corresponds to an *H* × *W* region with *C* immunofluorescence channels, and each channel contains a cellular marker (e.g., CD4^+^, CD8a^+^, CD19^+^, EpCAM^+^, IBA1^+^), capturing the spatial and phenotypic diversity of immune microenvironments.

Since patch-level labels {*y*_*ij*_} are unknown during the training process, the model operates under weak supervision, where only bag-level disease states *Y*_*i*_ are known. This formulation allows the network to learn discriminative representations by inferring which patches contribute most to the disease state label, despite substantial intra-bag heterogeneity across granulomas. Therefore, we adopt an attention-based Multiple Instance Learning (ABMIL) architecture to perform multi-class disease state classification. The model comprises three primary components: a patch encoder, an attention-based aggregator, and a classifier.

The patch encoder component *f*_*p*_(.;*θ*_*f*_) maps each patch *x*_*ij*_ into a high-dimensional embedding *D:*

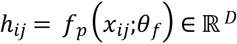

using a convolutional neural network (CNN) backbone trained to capture local morphological and cellular spatial features within each patch.

To aggregate patch embeddings into a single bag-level representation, we employ a gated attention mechanism that computes a weighted average over patch features:

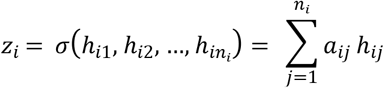

where *a*_*ij*_ denotes the normalized attention weight for the *j*^*th*^ patch. The weights form a probability distribution over all patches:

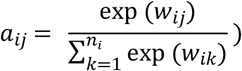

with the unnormalized attention logit defined as:

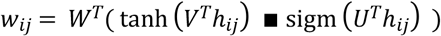

Here, *V, U* ∈ ℝ ^*D* × *L*^ are learned projection matrices, *W* ∈ ℝ ^*L* × 1^ is a learned vector, tanh, sigm denote hyperbolic tangent and sigmoid activations respectively, and ▪ represents element-wise multiplication. This gated attention mechanism allows the network to model complex, nonlinear dependencies between patches and its relevance, effectively emphasizing granuloma regions that are most informative for disease state discrimination.

The resulting aggregated embedding *z*_*i*_ ∈ ℝ ^*D*^ serves as a compact, disease-state-specific representation of the WSI, summarizing key morphological and immunological cues across all patches. Then, a fully connected layer with a SoftMax activation outputs a categorical probability distribution over the three disease states. The model is trained using categorical cross-entropy loss. We minimize this loss to enable the network to learn discriminative, attention-weighted feature representations that accurately capture the morphological and immunological signatures associated with distinct TB disease states.

### 3.3. Large Language Model-Based Interpretation of Spatial Patterns

The intricate spatial organization of immune cells within granulomas requires simultaneous integration of cell types, spatial distributions, expression intensities, and their interdependencies across disease states, demanding consideration of countless feature combinations across hundreds of tissue patches. To address this complexity, we employed a large language models (LLM) (31) to decode complex spatial patterns within high-attention regions of TB granulomas and identify recurring patterns across disease states (Figure 7).

We provided the model with comprehensive quantitative information for each granuloma patch, including: (1) predicted disease state (asymptomatic, chronic pulmonary TB, acute pulmonary TB), (2) cell types (CD19+, CD4+, CD8a+, EpCAM+, IBA1+), (3) spatial location of cells, (4) expression intensity values for each cell, (5) patch-level attention scores derived from the ABMIL model indicating discriminative capability, and (6) spatial coordinates defining patch positions within the granuloma structure. We tasked the model with identifying distinct cellular and spatial patterns that differentiate the three disease groups.

### 3.4. Implementation Details

For training the IF-UNet++ segmentation model, we employed five-fold cross-validation to ensure robust performance evaluation and model selection, using a batch size of 16 and a learning rate of 1×10^−3^. In each fold, 80% of the data was used for training, with 10% randomly selected from the training set for validation and the remaining 20% held out for testing. We monitored validation performance across all folds and selected the model with the best validation loss for granuloma segmentation for subsequent disease state classification, ensuring that the segmentation model generalized well to unseen data before being integrated into the downstream classification pipeline. For the ABMIL model, we employed five-fold cross-validation (70%/10%/20%) split with a learning rate of 1×10^−4^, and a batch size of 1.

## 4. Results

The overall goal of this study is to develop an end-to-end framework capable of segmenting granulomas from lung tissue sections, classifying TB disease states, utilization of LLMs for identifying cellular patterns, and spatial quantification of cellular composition in *M*.*tb*-infected DO mice. In this section, we present the performance of IF-UNet++, disease state classification, and an interpretation of cellular patterns using an LLM.

### 4.1. Granuloma Segmentation using IF-UNet++

Accurate delineation of granulomas is critical for extracting informative features and training reliable disease state classification models. Our segmentation network, IF-UNet++, effectively segmented granulomas, achieving an Intersection over Union (IoU) of 93.22% on the training set and 90.57% on the independent test set. These results highlight its strong generalization ability and robust segmentation performance. Representative examples of the segmentation output are shown in Figure 2. Panels A, B, and C illustrate automated granuloma segmentation across the three disease states: Progressor (survive < 60 days); asymptomatic infection, and controller (survive greater than 60 days) (3), with predicted granuloma boundaries outlined in yellow. The segmentation outputs exhibit high spatial concordance with granulomatous structures across all disease phenotypes, underscoring the model’s ability to accurately capture granulomas within complex pulmonary microenvironments. Panel D presents the confusion matrix demonstrating our framework’s classification performance, achieving high accuracy in distinguishing between the three TB disease states with minimal misclassification.

**Figure 2.**
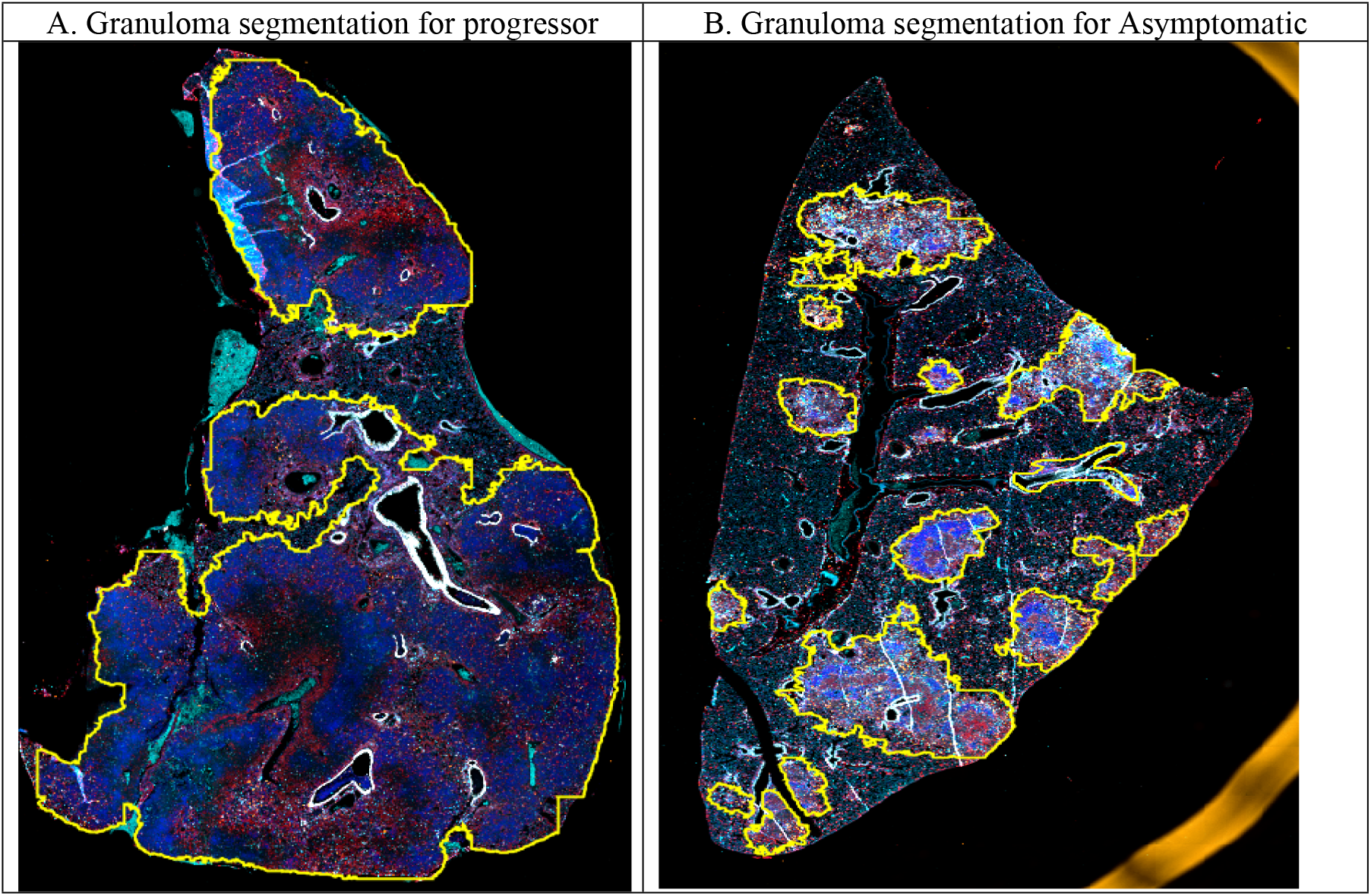
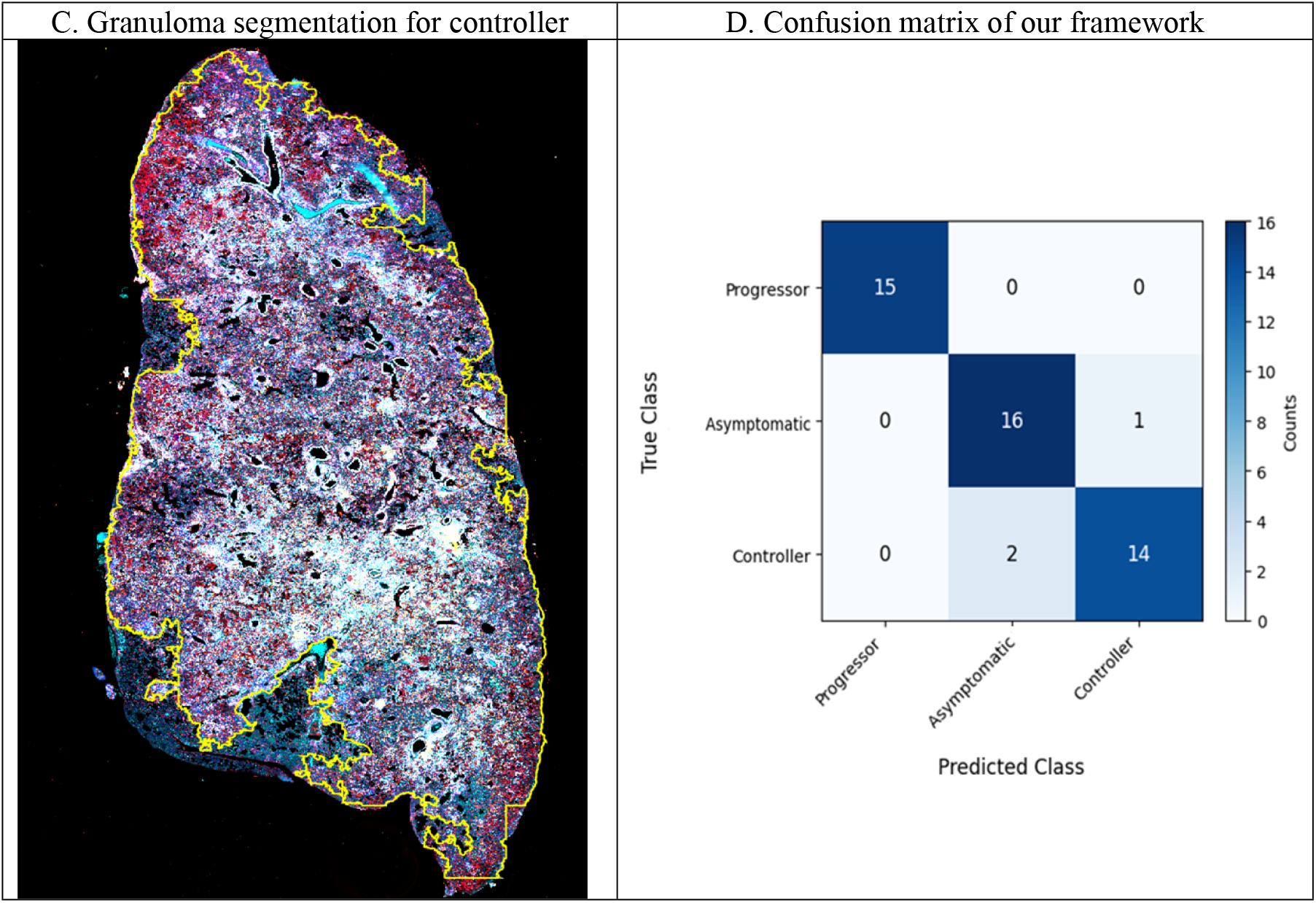
IF-UNet++ granuloma segmentation across disease states and classification performance. Representative examples of automated granuloma segmentation in lung tissue sections from DO mice across three disease states: **(A)** Progressor (acute pulmonary TB), **(B)** Asymptomatic (latent infection), and **(C)** Controller (chronic pulmonary TB). Each panel shows the original mIF image with predicted granuloma boundaries outlined in yellow. **(D)** Confusion matrix showing the classification performance of our framework in distinguishing between the three disease states.

### 4.2. Granuloma-Level Analysis of Immune Cell Populations

Statistical comparisons of immune cell abundance across disease states were performed using nonparametric tests because cell counts were non-normally distributed across granulomas. Group-wise differences were assessed followed by post hoc pairwise comparisons with appropriate multiple-testing correction. These analyses confirm that observed differences in immune cell composition across Progressor, Asymptomatic, and Controller groups reflect systematic biological variation rather than sampling variability.

To characterize the cellular heterogeneity underlying different TB disease states, we quantified the distribution of immune cell populations within each granuloma across the three disease groups (Figure 3). CD4+ T helper cells demonstrated a progressive increase in Asymptomatic in comparison with Progressor and with intermediate levels observed in Controllers (Figure 3A). CD8a+ cytotoxic T cells were most abundant in the Asymptomatic group compared to both the Progressor and Controller groups (Figure 3B). CD19+ B cells showed relatively sparse distribution across all disease states, with slightly elevated levels in Controllers (Figure 3C). EpCAM+ epithelial cell abundance was comparable across groups, suggesting similar cellular composition (Figure 3D). Notably, IBA1+ macrophages exhibited substantially higher macrophage density in Asymptomatic mice compared to both Progressor and Controller groups. When examining total cell abundance (Figure 4), Controller granulomas consistently exhibited the highest absolute cell counts across all immune cell types, including CD4+ T cells, CD8a+ T cells, CD19+ B cells, EpCAM+ epithelial cells, and IBA1+ macrophages.

**Figure 3.**
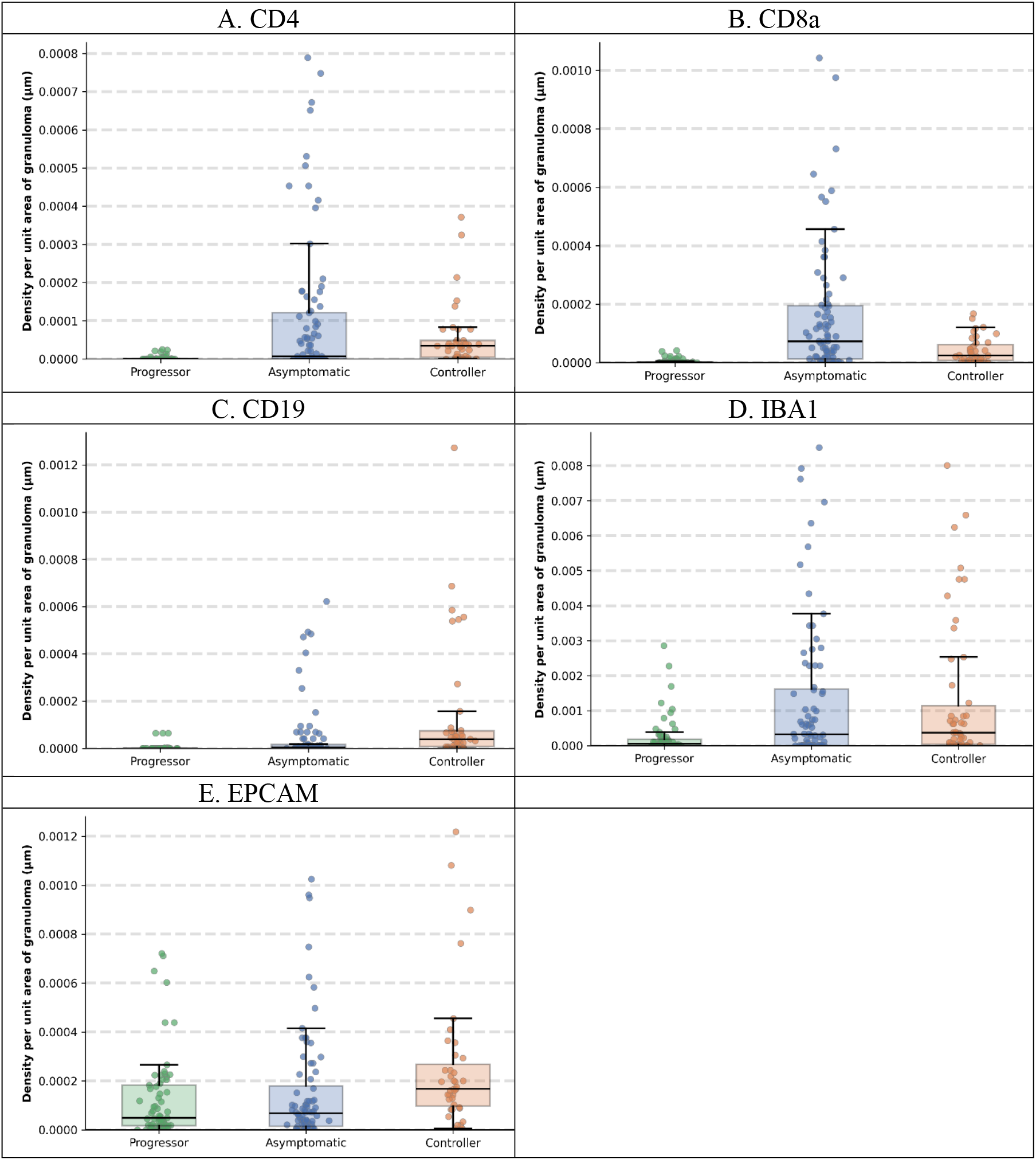
Density of immune cell populations per unit granuloma area across TB disease states. Box plots show density of cells, reported as density per unit granuloma area for **(A)** CD4+ T cells, **(B)** CD8a+ T cells, **(C)** CD19+ B cells, **(D)** IBA1+ macrophages, and **(E)** EpCAM+ epithelial cells across three disease groups: Progressor, Asymptomatic, and Controller. Each dot represents one granuloma; box plots show median, interquartile range, and whiskers extending to 1.5× IQR.

**Figure 4.**
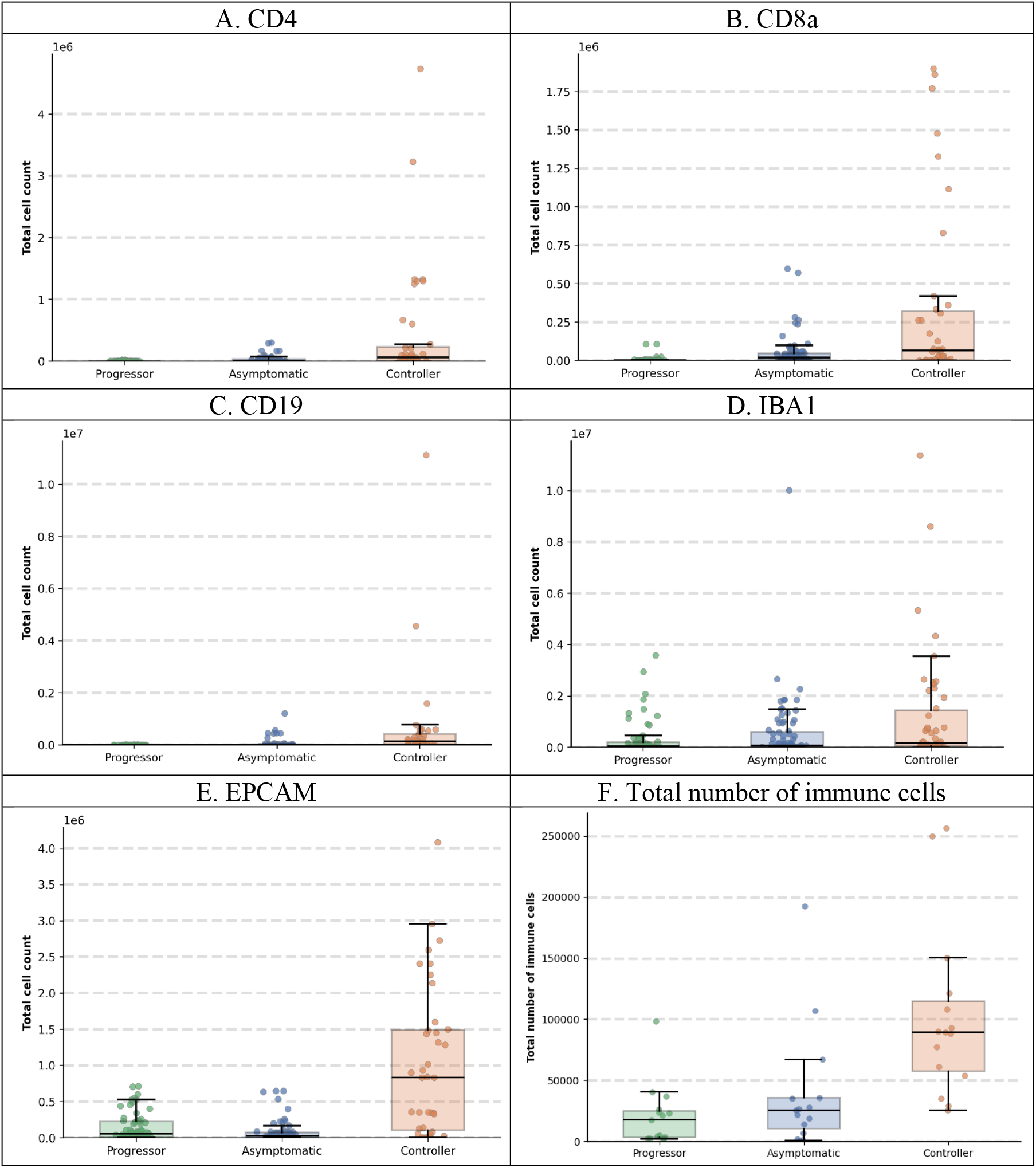
Total numbers of immune cells in granulomas across TB disease states. Box plots showing cell abundance for **(A)** CD4+ T cells, **(B)** CD8a+ T cells, **(C)** CD19+ B cells, **(D)** IBA1+ macrophages, and **(E)** EpCAM+ epithelial cells across three disease groups: Progressor, Asymptomatic TB, and Controller, **(F)** Total number of immune cells per lung tissue section. Each dot (4A-4E) represents one granuloma; box plots show median, interquartile range, and whiskers extending to 1.5× IQR.

### 4.4. Deep Learning Accurately Predicts Disease States From Granulomas

To differentiate between DO mice with asymptomatic *M*.*tb* infection, from progressors with acute pulmonary TB, and progressors with chronic pulmonary TB, we trained and evaluated our model on segmented granuloma regions. Our mIF-based classification model achieved an AUC of 0.982, with 90.0% accuracy, 91.1% sensitivity, and 95.1% specificity. In comparison, our previous H&E-based approach attained an AUC of 0.884, sensitivity of 71.9%, and specificity of 89.9% (Table 1). For multi-class evaluation, the reported AUC corresponds to a macro-averaged one-vs-rest formulation across the three disease states. These results demonstrate substantial performance improvements, particularly in sensitivity, highlighting the value of cell-type-specific spatial information provided by mIF imaging for disease state classification.

**Table 1.**
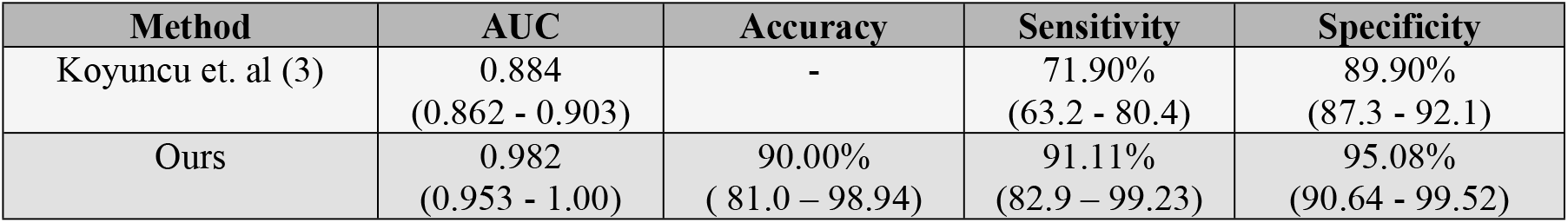
Performance comparison of disease state classification methods. Our model was evaluated using five-fold cross-validation to classify lung tissue sections into three disease states: asymptomatic infection, acute pulmonary TB, and chronic pulmonary TB. Values in parentheses represent 95% confidence intervals.

### 4.5. B-cells in Asymptomatic Infection Near Bronchiolar Epithelial Cells

To validate the peribronchiolar B-cell signature identified in previous study (3) and demonstrate the practical utility of our framework for spatial analysis, we quantified the distance between CD19+ B-cells and bronchiolar epithelial cells across disease states (Figure 5). Figure 5A presents a polar coordinate visualization of a representative granuloma from each disease state, illustrating the spatial distribution of immune cells as a function of distance from the center of the granuloma. Figure 5B shows Asymptomatic mice exhibit shorter distances between each bronchiolar epithelial cell and its nearest B-cell compared to both Progressors and Controllers, indicating closer spatial association and independently confirming the peribronchiolar B-cell signature (3) using mIF imaging and automated spatial analysis, validating previous observations through a complementary methodology.

**Figure 5.**
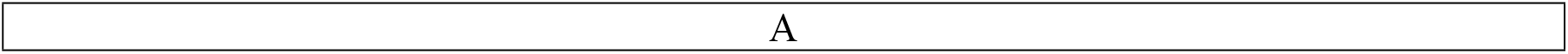
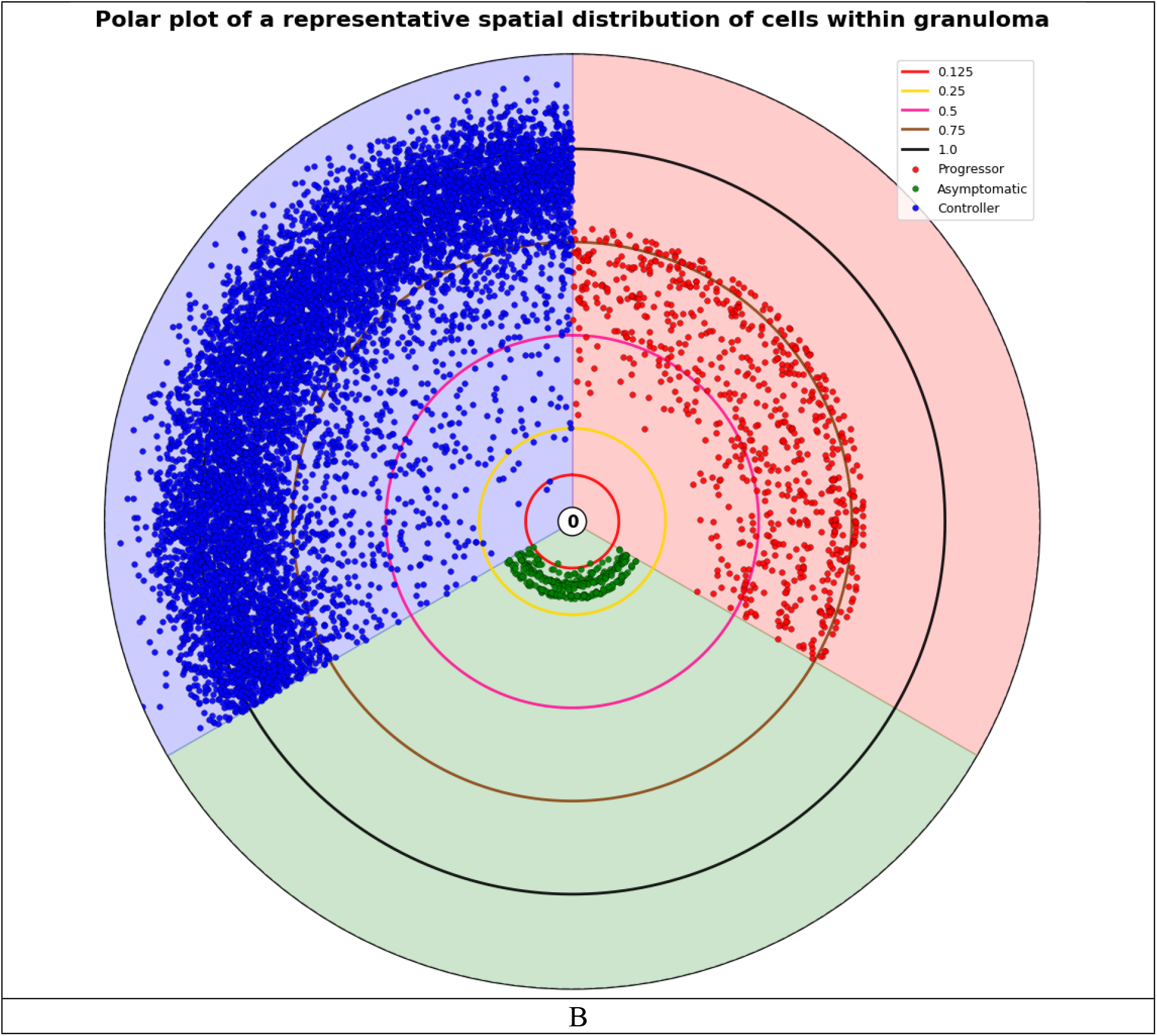
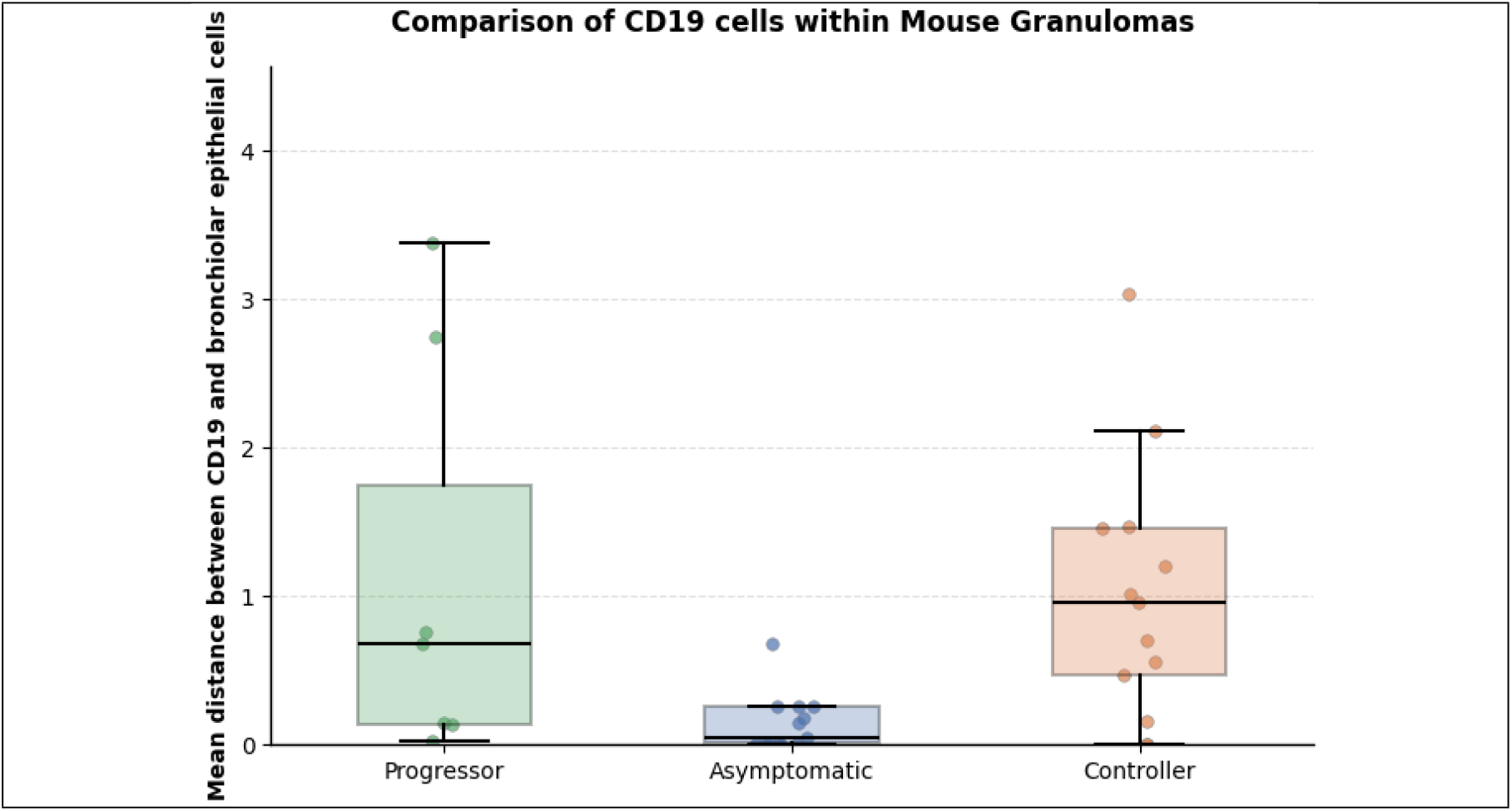
Spatial organization of B-cells in granulomas from *M*.*tb-*infected DO mice. (A) Polar plot of a representative granuloma showing spatial distribution of cells (Progressor: red, Asymptomatic: blue, Controller: green). Concentric circles indicate normalized radial distance from granuloma centroid. Each point represents the distance between each bronchiolar epithelial cell with nearest CD19+ B cell. (B) Mean distance between bronchiolar epithelial cells and nearest CD19+ B cell. Each point represents one mouse (averaged across all granulomas), demonstrating closest proximity in Asymptomatic.

### 4.6. Spatial Relationships Between T Cells and Macrophages Reveal Disease-Specific Immune Organization

To further characterize the spatial immune architecture within TB granulomas, we quantified the proximity between T cell populations (CD4+ and CD8a+) and IBA1+ macrophages across the three disease states (Figure 6). Figure 6A and 6B present the mean distances between CD4+ T cells and nearest macrophages, and CD8a+ T cells and nearest macrophages, respectively, across Asymptomatic, Controller, and Progressor groups. To assess whether T cells were positioned within functional proximity to macrophages, we defined a threshold of 80 μm radius as the spatial range for potential T cell-macrophage interactions based on (32). While both CD4+ and CD8a+ T cells showed variable distances to macrophages across individual mice, the overall proximity patterns differed among disease states with smallest mean distance between T cells and macrophages in asymptomatic mice in comparison with other groups. Figure 6C presents a two-dimensional visualization comparing the proportion of CD4+ cells within 80 μm of macrophages versus the proportion of CD8a+ cells within 80 μm of macrophages for each mouse. This analysis revealed distinct clustering patterns: Asymptomatic mice exhibited higher proportions of both T cell subsets in close proximity to macrophages, suggesting coordinated immune cell positioning that may facilitate effective pathogen control. Controllers displayed intermediate spatial patterns, while Progressors showed more dispersed and variable T cell-macrophage relationships, potentially reflecting impaired immune coordination. Figure 6D quantifies these observations, showing that Asymptomatic maintains the highest proportion of both CD4+ and CD8a+ T cells within 80 μm of macrophages, followed by Progressors with preferential CD4+ proximity and reduced CD8a+ proximity, and Controllers showing balanced but reduced overall T cell-macrophage proximity compared to Asymptomatic. These spatial profiling results demonstrate that the physical organization of T cells relative to macrophages differs systematically across TB disease states, with closer T cell-macrophage proximity in Asymptomatic potentially reflecting more effective immune surveillance and pathogen containment within granulomas.

**Figure 6.**
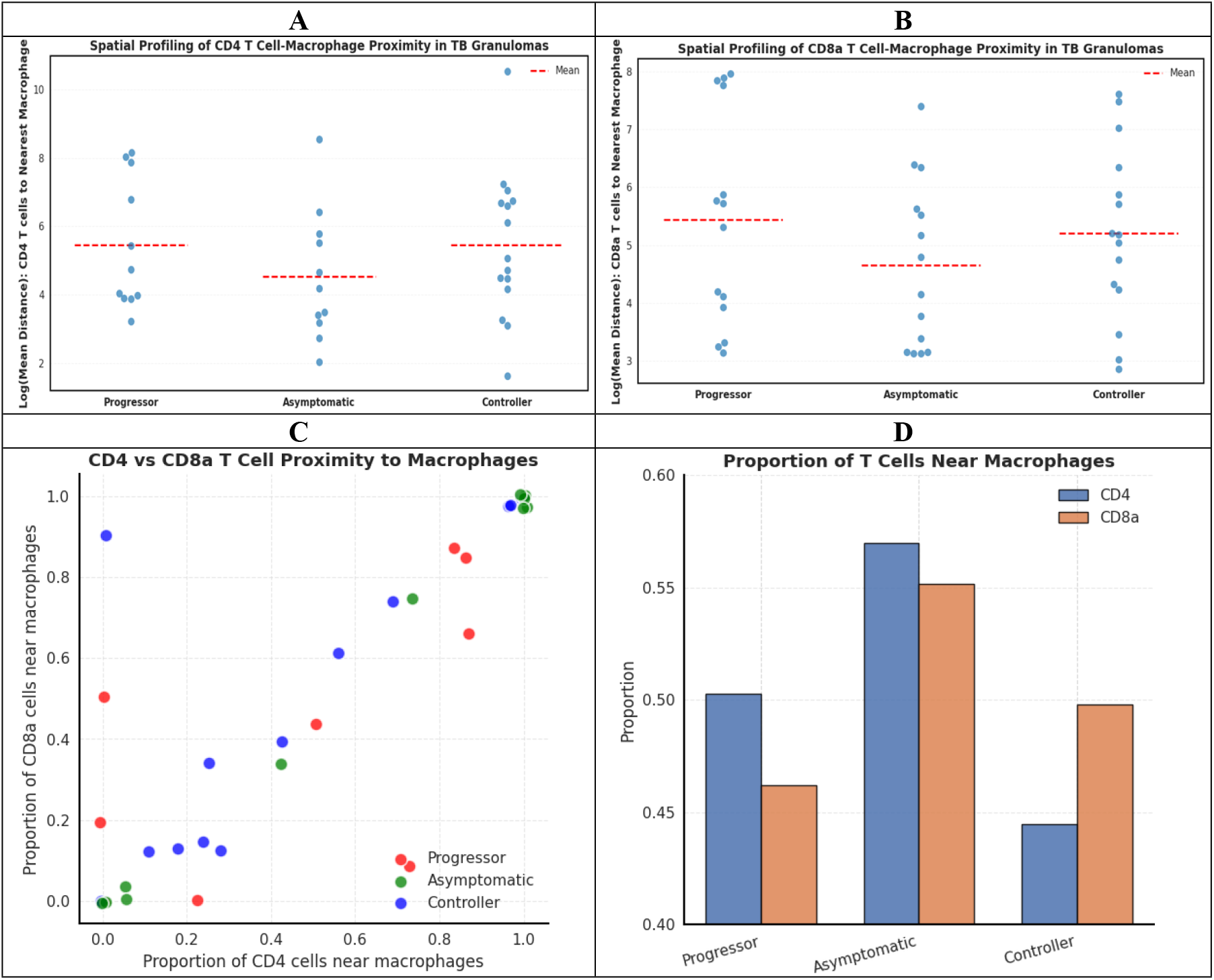
Spatial profiling of T cell-macrophage proximity in TB granulomas. (A) Swarm plot presenting a mean distance between CD4+ T cells and nearest IBA1+ macrophage per mouse. Each point represents one mouse (averaged across all granulomas); red dashed lines indicate mean values. (B) Swarm plot presenting mean distance between CD8a+ T cells and nearest IBA1+ macrophage. (C) Scatter plot showing joint T-cell proximity to macrophages. Each point represents one mouse, with x-axis showing the proportion of CD4+ T cells within 80 μm of macrophages and y-axis showing the proportion of CD8a+ T cells within 80 μm of macrophages. (D) Bar plot comparing the proportion of CD4+ and CD8a+ T cells within 80 μm radius of each macrophage across the three disease states, demonstrating differential spatial organization patterns characteristic of each TB disease phenotype.

**Figure 7.**
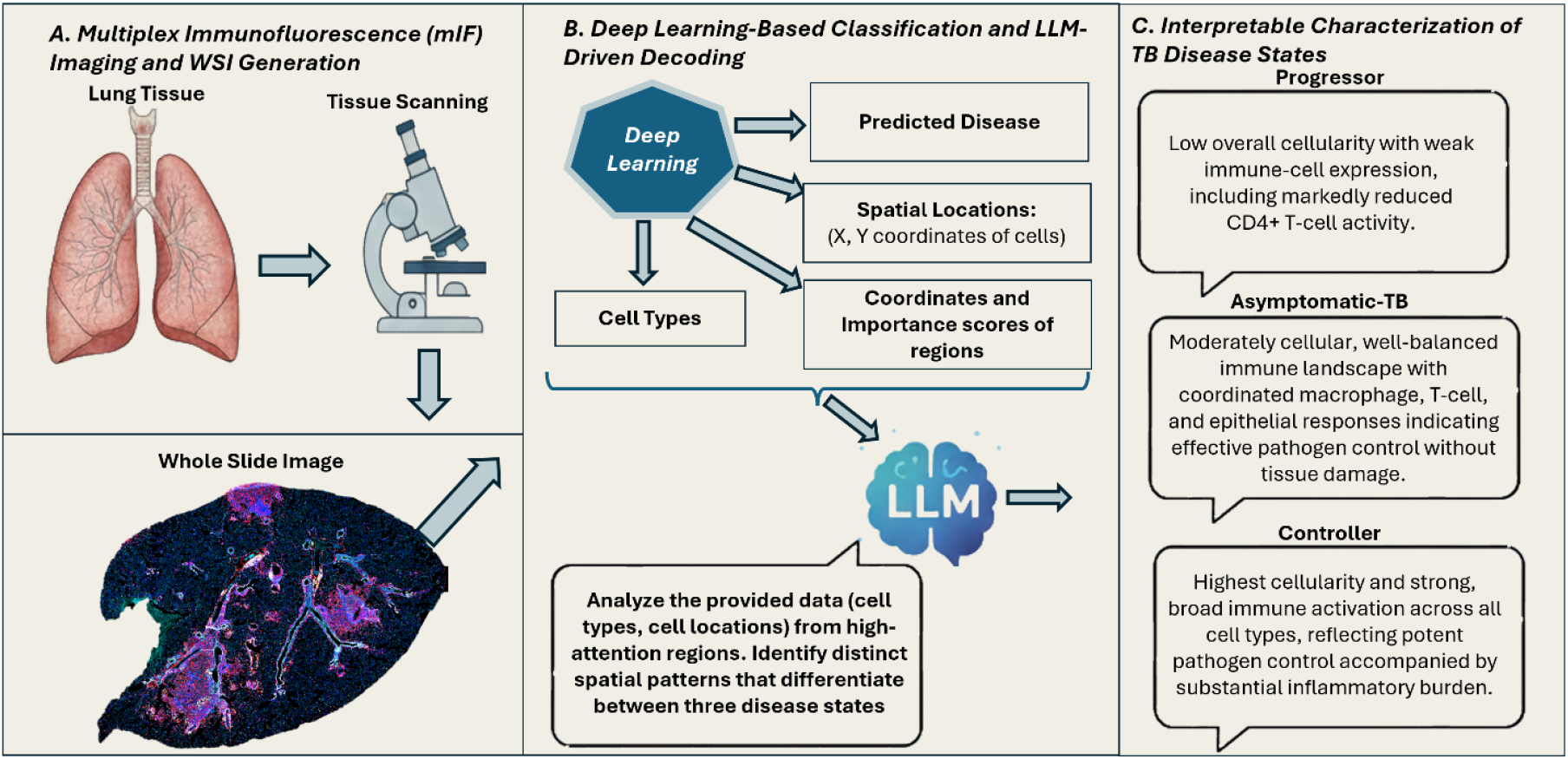
Framework for TB disease state classification and interpretation. **(A)** mIF imaging workflow from lung tissue to WSI generation. **(B)** Deep learning pipeline integrating cell type identification, spatial analysis, and LLM-based decoding to interpret disease-specific patterns. **(C)** Disease state characterizations: Progressor (low cellularity, weak immune response), Asymptomatic (balanced immune control), and Controller (high cellularity, strong inflammatory response).

### 4.7. Interpreting Disease-Specific Spatial Patterns using Large Language Models

Multi-modal large language models have demonstrated promising capabilities for analyzing histopathology images [33-35]. Recent approaches [33-34] enable automated quantification and interpretation of morphological markers in large data cohorts, addressing key limitations of manual quantification, which is time-consuming and subject to interobserver variability. However, these methods remain limited to H&E images. Multi-modal large language models have demonstrated promising capabilities for analyzing histopathology images (33-35). Recent approaches (33, 34) enable automated quantification and interpretation of morphological markers in large data cohorts, addressing key limitations of manual quantification, which is time-consuming and subject to interobserver variability. Here, we leverage a LLM as an integrative reasoning engine to synthesize multi-modal computational outputs (cell identities, spatial coordinates, expression intensities, and attention scores) into interpretable biological narratives, enabling an automated hypothesis-free pattern discovery across complex spatial immune phenotypes. Using this approach, LLMs (31, 36, 37) serves as a higher-level interpretive layer that summarizes recurring, disease-specific immune architectures across high-impact granuloma regions. We tasked the model with identifying distinct cellular and spatial patterns that differentiate the three disease groups.

The following are the findings of the LLM:

1. Progressors are characterized by relatively low overall cellularity and dysfunctional expression of immune cells. While CD4+ T cells are numerically present, their expression is notably low.
2. Asymptomatic displays a balanced cellular landscape with moderate overall cellularity and expression. A well-coordinated expression of macrophages, CD4+ and CD8+ T cells, and epithelial cells. This immune response indicates effective pathogen containment without inducing excessive inflammation or tissue damage, representing an optimal host-pathogen equilibrium.
3. Controllers exhibit the highest overall cellularity and widespread, intense immune expression level across all cell types. High expression of CD4+ and CD8+ T cells, B cells, and macrophages signifies a robust, hyper-inflammatory response. This aggressive, broad-spectrum immune mobilization effectively contains the pathogen but likely contributes to significant immunopathology due to excessive inflammation and tissue damage.

Importantly, these interpretations were derived from model-generated quantitative inputs rather than raw image data. The identified patterns were concordant with independently quantified differences in cellular abundance and spatial proximity, supporting their biological plausibility.

## 5. Discussions and Conclusions

This study presents an end-to-end framework for automated analysis of lung granulomas from *M*.*tb*-infected DO mice using mIF images stained for CD4 and CD8a T cell subsets, B cells, macrophages, and bronchiolar epithelial cells. Our framework involves automated granuloma segmentation, disease-state classification, and interpretability analysis to decode the organization of immune cells within TB granulomas, addressing fundamental limitations of manual annotation and traditional H&E-based computational approaches.

Our proposed IF-UNet++ segmentation model achieved robust performance in delineating granulomatous regions, with an IoU of 93.22% on the training set and 90.57% on the independent test set. These results demonstrate strong generalization and establish accurate spatial boundaries, which are essential for downstream feature extraction and disease-state classification (Figure 2). Building on these segmentation outputs, our attention-based multiple instance learning (ABMIL) framework achieved substantial improvements in disease-state classification over previous approaches. Compared to our previous H&E-based method, the mIF-based model demonstrated an 11.1% increase in AUC (0.982 vs. 0.884), a 26.7% increase in sensitivity (91.1% vs. 71.9%), and a 5.8% increase in specificity (95.1% vs. 89.9%). These marked performance gains, particularly the substantial enhancement in sensitivity, highlight the value of cell-type-specific spatial information provided by mIF images.

Our spatial and cellular analyses revealed distinct patterns that differentiate TB disease states in DO mice at the granuloma level. When examining density per granuloma area (Figure 3), we observed that CD4+ T cells and CD8a+ T cells were most abundant in Asymptomatic, suggesting a balanced adaptive immune response associated with effective pathogen control. In contrast, IBA1+ macrophages showed substantially higher density in Asymptomatic mice compared to both the Progressor and Controller groups, indicating a well-regulated innate immune response. Analysis of total cell counts (Figure 4) revealed that Controller granulomas consistently exhibited the highest absolute cell numbers across all immune cell types. Critically, our analysis also validated that the peribronchiolar B-cell signature identified in our previous studies (3), where asymptomatic mice demonstrated significantly closer proximity between CD19+ B-cells and bronchiolar epithelial cells compared to Progressor and Controller groups (Figure 5). This independent confirmation through mIF imaging and automated spatial quantification not only validates prior findings but also demonstrates the translational utility of our framework for discovering and validating spatial biomarkers that are associated with protective immunity versus disease progression. These cell-type-specific spatial patterns may provide mechanistic insights into the differential immune responses underlying asymptomatic infection, acute pulmonary TB, and chronic pulmonary TB.

Granulomas organization, the cell types within the granulomas, and the function of immune cells within granulomas and their subregions are important to understanding clinical cases of pulmonary T in humans and in animal models of *M*.*tb* infection (38-42). However, few studies compare immune cell types across different disease states. Here, our spatial profiling analysis quantifies density per unit area of granuloma, total count per lung tissue section, and distances of immune cells in lung granulomas from DO mice with acute pulmonary TB, asymptomatic infection, and chronic pulmonary TB. Our work reveals that T cell-macrophage proximity patterns are disease-state specific and may serve as functional indicators of immune coordination. Our findings that asymptomatic mice maintain the closest spatial relationships between both CD4+ and CD8a+ T cells and IBA1+ macrophages, with the highest proportions of both T cell subsets positioned within the 80 μm range. This spatial organization may facilitate effective and coordinated pathogen containment mechanisms critical for maintaining control of asymptomatic latent infection. In contrast, progressors exhibited more dispersed and variable T cell-macrophage spatial relationships, potentially reflecting breakdown of organized immune architecture and impaired cell-cell communication necessary for granuloma function. Controllers displayed intermediate patterns, suggesting partially maintained but suboptimal immune coordination. These findings extend beyond simple cell abundance metrics to reveal that the physical organization of immune cells within granulomas is a critical determinant of disease outcome, highlighting the importance of spatial profiling approaches for understanding TB pathogenesis and identifying potential therapeutic targets aimed at restoring functional immune architecture.

Another novel contribution of this work is the integration of LLMs to interpret complex immune organization within TB granulomas. We provided the LLM with comprehensive multi-modal data, including predicted disease states, cell identities, spatial coordinates, protein expression intensities, and patch-level attention scores derived from the model. Through this integrative analysis, the LLM identified disease-specific spatial patterns that were independently validated by our quantitative analyses. For example, the LLM identified coordinated spatial clustering of macrophages, CD4+ T cells, and CD8+ T cells in asymptomatic mice, which was corroborated by the spatial relationship analyses shown in Figures 6A and 6B. Similarly, the LLM’s identification of elevated overall cellularity in controllers was validated by the quantitative measurements presented in Figure 4. These findings provide an automated, scalable framework for discovering distinctive spatial immune patterns within granulomas without requiring pre-defined hypotheses or manual feature selection.

The translational impact of this work is fundamentally enabled by our use of the DO mouse population, which directly addresses a critical barrier in TB research: the failure of traditional inbred mouse models to recapitulate the heterogeneity of human disease. Unlike inbred strains, DO mice capture the broad spectrum of clinical outcomes seen in human populations from LTBI to progressive pulmonary disease, thereby providing an unprecedented animal model for training computational models on biologically realistic disease variability. This genetic diversity not only strengthens the translational relevance of our findings but also enables our framework to learn from granuloma phenotypes that mirror the complex host-pathogen interactions underlying human TB pathogenesis, making our identified spatial biomarkers directly relevant to human disease heterogeneity.

Beyond its application to murine models, the proposed framework is directly extensible to emerging human spatial immunology datasets, including multiplex immunofluorescence and spatial transcriptomics of lung tissue. By explicitly modeling cell-type–specific spatial relationships within granulomas, this approach provides a scalable pathway for translating insights from genetically diverse animal models to human tuberculosis, where similar immune architectures underlie heterogeneous clinical outcomes.

Despite several breakthroughs in this study, there are several limitations. First, our dataset included only 48 mIF-stained lung tissue sections. Although this dataset provided a strong foundation, expanding to larger, multi-institutional cohorts will be essential to fully assess generalizability and clinical applicability. Second, our interpretability analysis focused on high-attention patches identified by the ABMIL model heavily relies on a user-defined hyperparameter governing the number of selected regisons, ‘K.’ While this strategy prioritizes biologically salient tissue compartments, future work will explore adaptive selection strategies and whole-granuloma reasoning to reduce parameter sensitivity further. Third, while the LLM-based interpretation provides insights into cellular patterns within high-attention regions, this approach has inherent limitations. The LLM operates as a black-box system, making it difficult to fully understand how it integrates spatial coordinates, cell identities, and expression values to generate its interpretations. Given these constraints and the potential for AI-generated misinterpretations, maintaining pathologist oversight remains essential to validate LLM outputs and ensure biological accuracy. Additionally, due to computational constraints, the LLM analyzes only high-attention patches rather than entire granulomas, which may not capture the complete organization of immune cells or address broader biological phenomena occurring at the whole-granuloma or whole-lung level. Furthermore, the LLM’s reasoning is inherently constrained by the specific set of biomarkers included in the study panel, and the patterns it identifies might not always align with or capture the full spectrum of human-identified biological patterns. This discordance may arise because relevant cellular populations or biological processes that are recognizable through other markers or visual morphological features may not be adequately represented in our current marker panel, and hypothesis-free pattern discovery, potentially limiting the LLM’s ability to detect or interpret certain biologically significant phenomena. For instance, while progressors are known to exhibit large neutrophilic infiltrates, the absence of a neutrophil-specific marker in our panel means that the LLM’s interpretations cannot directly assess this phenomenon, and conclusions are necessarily limited to the cellular populations and markers available in the dataset. Finally, the MIL framework assumes patch independence and may not fully capture long-range spatial dependencies during training; integrating graph-based or transformer architectures in future work could enhance spatial reasoning capabilities.

In summary, this study establishes a foundation for automated, interpretable, and biologically grounded characterization of TB granulomas. Our proposed framework achieves precise delineation of TB granulomas, accurate disease-state classification, and interpretation of complex spatial cellular patterns. The substantial improvement over traditional H&E-based methods highlight the advantage of incorporating cell-type–specific spatial information into computational pathology workflows.

## Contributions

U.S. performed data preprocessing, experimental design, validation, and manuscript writing. G.B., M.N.G., and M.K.K.N. provided feedback on study design, validation, and manuscript editing. M.N.G., contributed to validating the experimental design, and AI evaluation. N.A.C. contributed to clinical assessment, and editing the manuscript. G.B. contributed to slide annotation and clinical assessment. G.B. and M.K.K.N. conceptualized and designed the study and supervised the research.

## Acknowledgements

We thank Dr. Sam Telford III and the biocontainment staff at the New England Regional Biosafety Laboratory at Tufts University Cummings School of Veterinary Medicine, North Grafton, MA, USA, for support with biosafety level three facilities. We thank the NEIDL Comparative Pathology Laboratory (NCPL) for their support generating the mIF images. The authors gratefully acknowledge the Ohio Supercomputer Center for providing high-performance computing resources under its contract with The Ohio State University College of Medicine. The instruments utilized for IHC and fluorescent whole slide scanning were acquired with support from NIH SIG grants (S10OD026983 and S10OD030269).

